# CRY-BARs: Versatile light-gated molecular tools for the remodeling of membrane architectures

**DOI:** 10.1101/2022.01.28.478241

**Authors:** Anna I. Wurz, Wyatt Paul Bunner, Erzsebet M. Szatmari, Robert M. Hughes

## Abstract

BAR (Bin, Amphiphysin and Rvs) protein domains are responsible for the generation of membrane curvature and represent a critical mechanical component of cellular functions. Thus, BAR domains have great potential as components of membrane-remodeling tools for cell biologists. In this work, we describe the design and implementation of a family of versatile light-gated I-BAR domain containing tools (‘CRY-BARs’) with applications in the remodeling of membrane architectures and the control of cellular dynamics. By taking advantage of the intrinsic membrane binding propensity of the I-BAR domain, CRY-BARs can be used for spatial and temporal control of cellular processes that require induction of membrane protrusions. Using cell lines and primary neuron cultures, we demonstrate that the CRY-BAR optogenetic tool reports membrane dynamic changes associated with cellular activity. Moreover, we provide evidence that Ezrin acts as a relay between the plasma membrane and the actin cytoskeleton and therefore is an important mediator of switch function. Overall, CRY-BARs hold promise as a useful addition to the optogenetic toolkit to study membrane remodeling in live cells.

## INTRODUCTION

Membrane bound architectures, including filopodia, lamellipodia, and dendritic spines in neurons, are critical for a cell’s ability to transmit and respond to extracellular cues. As such, chemists and biologists have developed numerous optogenetic and chemo-optogenetic tools to control cellular architectures via manipulation of plasma membrane dynamics (Kichuk, Carrasco-López, & Avalos, 2021; Klewer & Wu, 2019; Ueda & Sato, 2018). However, these tools have to date incorporated a relatively small fraction of the numerous proteins involved in the building and dismantling of these architectural features. In particular, non-enzymatic proteins have been overlooked: while numerous optogenetic strategies exist for enzyme-mediated control of membrane dynamics (Khamo, Krishnamurthy, Chen, Diao, & Zhang, 2019; C.P. O’Banion et al., 2017; Colin P O’Banion, Vickerman, Haar, & Lawrence, 2019; Shaaya et al., 2020; Wu et al., 2009; Wu, Wang, He, Montell, & Hahn, 2011), far fewer have incorporated non-enzymatic proteins as the basis of optogenetic switch action (Redchuk et al., 2020). As mechanical control of membrane architecture by non-enzymatic proteins is a critical component of cellular signaling (Hughes & Kumar, 2016), addressing this oversight will be a bridge to a more comprehensive understanding of cellular dynamics and function.

In recent years, the critical role of proteins involved in membrane curvature inducing and sensing has come to light (Antonny, 2011). In particular, BAR domain containing proteins possess diverse activities at the plasma membrane (Linkner et al., 2014; Prévost et al., 2015; Pykäläinen et al., 2011; Saarikangas et al., 2015, 2011; Yu et al., 2011; Zhao, Pykäläinen, & Lappalainen, 2011); as such, a heightened understanding of these roles has invited recent investigations of their suitability for control with optogenetic tools (Jones, Liu, & Cui, 2020). In this work, we investigate the potential of an inverse BAR (I-BAR) domain from the MTSS1 protein (Missing in Metastasis 1) to serve as an actuator of membrane architecture and plasma membrane dynamics. We demonstrate that the I-BAR domain from MTSS1, in conjunction with the Cry2 optogenetic switch (Kennedy et al., 2010), is the basis of a versatile optogenetic approach (‘CRY-BAR’) for controlling membrane dynamics and cellular architecture. We also provide insight into the mode of action of CRY-BAR by showing that Ezrin, a membrane and cytoskeletal relay protein, is linked to CRY-BAR’s ability to induce light-activated membrane remodeling and spatial restriction of cellular dynamics.

## MATERIALS AND METHODS

### Plasmids and Cloning

Cloning of IBAR-Cry2-mCh, Cry2-mCh-IBAR, IBAR-Cry2-mCh-WH2, and Cry2-mCh-MTSS1 constructs was conducted using a previously described cloning scheme (Salem et al., 2020). Briefly, the IBAR domain from MTSS1, the WH2 domain from MTSS1, and MTSS1 were PCR amplified from human MTSS1 cDNA obtained from the Arizona State University DNA repository (DNASU ID HsCD00746054). N-terminal IBAR genes were cloned into Cry2PHR-mCherry (Addgene #26866) using NheI and XhoI restriction sites (Kennedy et al., 2010). C-terminal IBAR, WH2, and MTSS1 genes were cloned into Cry2PHR-mCherry using BsrgI and NotI restriction sites. Ezrin-GFP was expressed from plasmid pHJ421 (Addgene #20680) (Hao et al., 2009). Midi prep quantities of DNA of each construct were created from *E. coli* and collected for cell transfection.

### Cell lines and Transfection

HEK293T cells (passage 8) used for these experiments were purchased from ATCC and were cultured in DMEM medium containing 10% FBS and 1% Penicillin-Streptomycin. Cultures were transfected at 70% confluency with the Lipofectamine 3000 reagent (Invitrogen) following manufacturer’s suggested protocols. Briefly, for dual transfections in 35 mm glass bottom dishes for cell imaging or 6-well culture plates for lysis, plasmid DNA was combined in a 1:1 ratio (1,250 ng per plasmid) in 125 µl of Opti-Mem, followed by the addition of 5 µl of P3000 reagent. For single transfections, 2,500 ng of plasmid DNA was used per transfection. In a separate vial, 3.75 µl of Lipofectamine 3000 were added to 125 µl of Opti-Mem. The two 125 µl solutions were combined and allowed to incubate at room temperature for 10 min, followed by dropwise addition to cell culture. Transfection solutions remained on cells overnight.

Transfected cells were maintained at 37°C and 5% CO_2_ in a humidified tissue culture incubator, in culture medium consisting of DMEM supplemented with 10% FBS and 1% Penicillin-Streptomycin.

### Neuron cultures and Transfection

Postnatal dissociated cortical neuron cultures were prepared as previously described (Bunner, Dodson, Szatmari, & Hughes, 2021; Chang et al., 2017), from newborn B6 mice. Neurons were plated into 35 mm glass bottom Petri dishes at a 1 million/ml density in culture medium consisting of BME supplemented with 10% BCS and 1% Penicillin-Streptomycin. On day in vitro 2 (DIV2), culture medium was changed to Neurobasal A medium, supplemented with B27-plus reagent (Invitrogen), Glutamax and 1% Penicillin-Streptomycin. Neurons were transfected with the CryBAR optogenetic system (6 µg plasmid/plate) on DIV5 using Lipofectamine LTX reagent (Invitrogen). On DIV7, culture medium was removed and neurons were placed in imaging solution (Mg-free HEPES buffered aCSF (Sun, Smirnov, Kamasawa, & Yasuda, 2021)) Live cell imaging was performed before and after illumination using a Leica DMi8 Live Cell Imaging System.

### Fixed and Live-cell Imaging Preparation

#### Fixed cell experiments

Immediately prior to fixation, transfected HEK293T cells were either kept in dark conditions or continually illuminated with LED blue light (Sunlite LED Par30 Reflector, Item #80021, 4 Watts, 120 Volt, 66.21 µmol/s/m^2^; placed 10 cm from cell culture dishes) for 5 min. Cells were washed with Dulbecco’s PBS (with calcium and magnesium; 1x 1 mL), then fixed for 10 min at room temperature with pre-warmed 4% Paraformaldehyde solution in DPBS (37°C; prepared from 16% PFA (Electron Microscopy Sciences)). Following fixation, cells were washed with DPBS, then stored in DPBS at 4°C.

#### Live cell experiments

Transfected HEK293T cell media was replaced with 10% FBS in Leibovitz’s L-15 Medium. Cells were allowed to equilibrate in the live cell incubation system (OKOLab) for 10 minutes prior to beginning the illumination sequence.

### Imaging

#### Confocal Microscopy

Confocal images of fixed cells were collected with a Zeiss LSM 700 laser scanning microscope using ZEN Black 2012 software. Fluorescence images were colorized and overlaid using FIJI software.

#### Widefield Microscopy

A Leica DMi8 Live Cell Imaging System, equipped with an OKOLab stage-top live cell incubation system, LASX software, Leica HCX PL APO 63x/1.40-0.60na oil objective, Lumencor LED light engine, CTRadvanced+ power supply, and a Leica DFC900 GT camera, was used to acquire images. Exposure times were set at 200 ms (mCherry, 550 nm) and 50 ms (GFP, 480 nm), with LED light sources at 50% power, and images acquired every 30 seconds over specified time course.

#### Western blotting

Transfected HEK293T cells were lysed with 200 µL of M-PER lysis buffer (Thermo Scientific) containing 1X Halt protease-phosphatase inhibitor cocktail (Thermo Scientific). After 10 min on a rotary shaker at room temperature, lysates were collected and centrifuged for 15 min (94 rcf; 4°C). The supernatants were combined with Laemmli SDS sample buffer (Alfa Aesar) and incubated at 65°C for 10 min. The resulting lysates were subjected to electrophoresis on a 10% SDS-PAGE gel and then transferred onto PVDF membranes (20 V, 800 min). Membranes were then blocked for 1 h with 5% BSA in 1x TBS with 1% Tween (TBST), followed by incubation with primary antibody (Anti-mCherry antibody (Cell Signaling) 1:1000 dilution in 5% BSA – TBST; Anti-GAPDH antibody (Invitrogen) 1:1000 dilution in 5% BSA – TBST) overnight at 4°C on a platform rocker. The membranes were then washed 3x for 5 min each with TBST and incubated with the appropriate secondary antibody in 5% BSA – TBST (1 h; room temperature). After washing 3x for 5 min with TBST, the membranes were exposed to a chemiluminescent substrate for 5 min and imaged with an Azure cSeries imaging station.

## Results and Discussion

Increased MTSS-1 activity has been associated with exercise-induced enhancement of synaptic function (Chatzi et al., 2019). This pro-synaptic plasticity effect of MTSS-1 is attributed to the presence of an I-BAR domain within its structure. Moreover, I-BAR domains have also been shown to promote formation and stabilization of dendritic spines (**Fig. 1A**), leading to improved synaptic function and resilience against neurodegeneration (Saarikangas et al., 2015). To gain insights into the molecular mechanisms that control membrane expansion associated with cell morphology changes, we incorporated the I-BAR domain from MTSS-1 into a light-activated optogenetic protein (Cry2PHR; **Fig. 1B-C**), creating a light-activatable I-BAR protein (CRY-BAR). For this investigation, we engineered four permutations of the CRY-BAR switch: IBAR-Cry2-mCh, IBAR-Cry2-mCh-WH2, Cry2-mCh-IBAR, and Cry2-mCh-MTSS1 (**Fig. 1C**). Each construct was confirmed by Sanger and NextGen sequencing and expressed at the expected molecular weight (**Supporting Fig. 1**).

**Figure 1.**
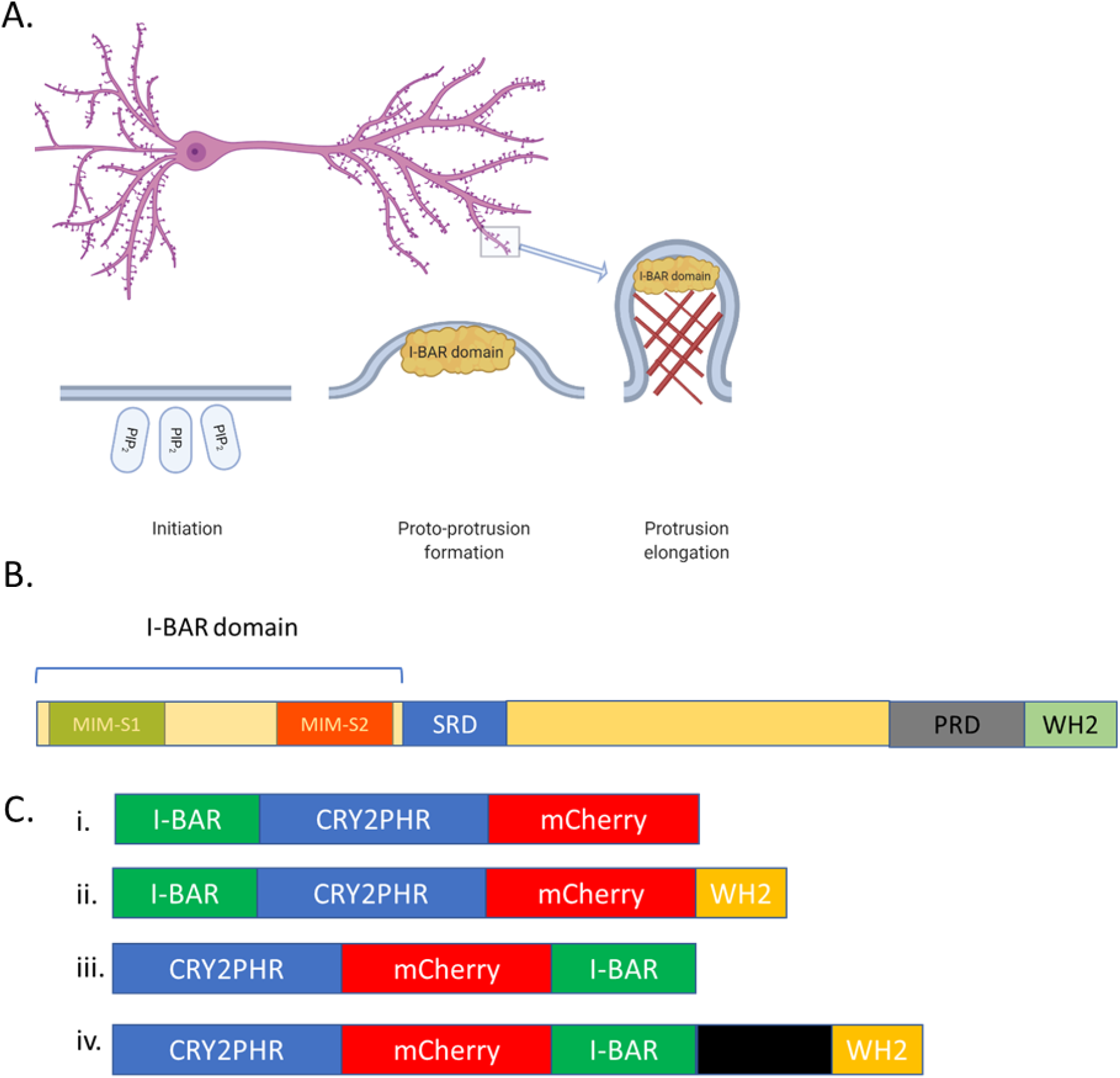
BAR-mediated initiation of dendritic spine generation in neurons. A.Phosphoinositide (PIP2) signaling recruits I-BAR domain proteins to the plasma membrane, inducing a proto-protrusion. Subsequent recruitment of actin (red crosshatches) and actin binding and remodeling proteins promotes protrusion elongation. Mature dendritic spines (purple nodules on dendrite) are mushroom-shaped bulbous protrusions and major sites of excitatory synaptic transmission in the mammalian brain. B. Diagram of MTSS-1, an I-BAR domain containing protein. The N-terminal I-BAR domain (250 amino acids) is comprised of three alpha helices. MIM-S1 and MIM-S2 represent the I-BAR domain dimerization interface as determined from X-ray crystallography [34]. SRD = serine-rich domain; PRD = proline-rich domain; WH2 = Wiskott - Aldrich syndrome homology region. C. Schematic of CRY-BAR optogenetic switches: i. The N-terminal I-BAR domain (250 amino acids) is fused to photoreceptor protein CRY2, which is fused to the red fluorescent mCherry protein for visualization purposed. ii. At the C-terminus, proline-rich and WH2 domains can be included for actin-recruitment. iii. The N-terminal I-BAR domain is fused to the C-terminus of mCherry. iv. The intact MTSS1 protein is fused to the C-terminus of mCherry. Graphic in panel A generated with BioRender (https://biorender.com/).

CRY-BAR constructs were initially probed for their response to blue light in the presence of CIB-CAAX, a membrane localized binding partner to Cry2, in HEK293T cells. As anticipated, light activation of the Cry2-mCh control resulted in rapid recruitment to the plasma membrane (**Fig. 2A** and **3A**). By contrast, the I-BAR domain containing constructs exhibited less apparent light-activated recruitment to the plasma membrane (**Fig. 2A, 2C**, and **3B**). We attributed this to the PIP2-binding propensity of the I-BAR domain, which results in significant pre-localization of the CRY-BAR constructs (readily observed in the pre-illumination images in **Fig. 2A** and **2C**). Due to the significant pre-localization of these constructs, we then investigated whether they might exhibit light activation in the absence of CIB-CAAX. These experiments revealed significant light activated clustering/membrane recruitment (**Fig. 2B and 2D**) in the absence of CIB-CAAX. This effect was observed for all of the I-BAR containing constructs with the exception of Cry2-mCh-MTSS1, which accumulated primarily in non-light responsive clusters. As a result, this construct was not pursued further.

**Figure 2.**
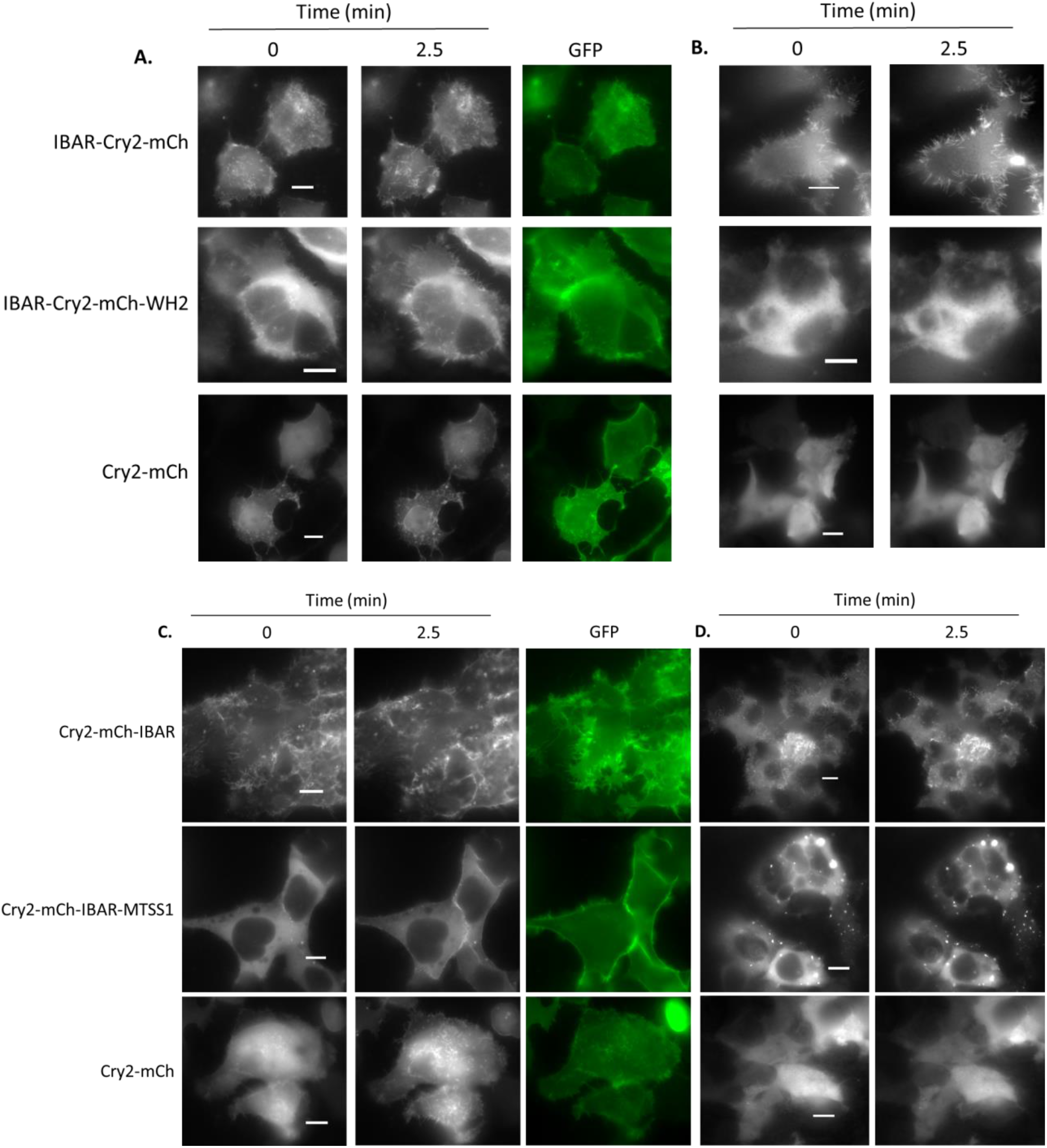
Light-activated responses of CRY-BAR constructs. A. HEK293T cells were co-transfected with CIB-CAAX and CRY-BAR fusions IBAR-Cry2-mCh and IBAR-Cry2-mCh-WH2, respectively, and imaged on a widefield fluorescent microscope. B. HEK293T cells were transfected with CRY-BAR fusions IBAR-Cry2-mCh and IBAR-Cry2-mCh-WH2, respectively and imaged on a widefield fluorescent microscope. C. HEK293T cells were co-transfected with CIB-CAAX and CRY-BAR fusions Cry2-mCh-IBAR and Cry2-mCh-MTSS1, respectively, and imaged on a widefield fluorescent microscope. D. HEK293T cells were transfected with CRY-BAR fusions Cry2-mCh-IBAR and Cry2-mCh-MTSS1, respectively and imaged on a widefield fluorescent microscope. Pre = image before 480 nm light exposure; Post = image after 480 nm light exposure. Images acquired every 30 s. Scale bars = 10 microns.

**Figure 3.**
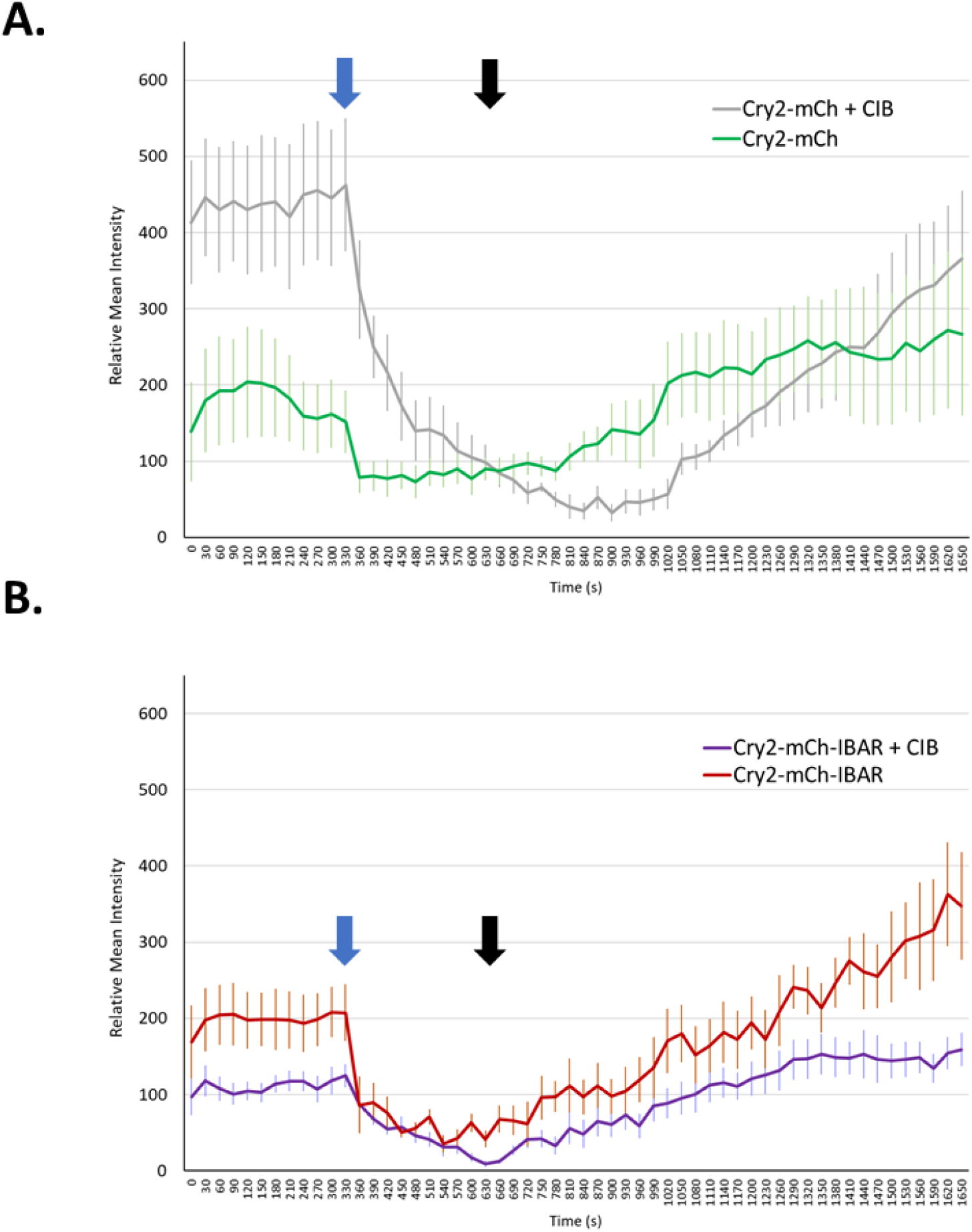
Analysis of Light-activated responses of CRY-BAR constructs. A. Cry2-mCherry construct shows large decrease in cytosolic fluorescence intensity in the presence of plasma membrane anchored CIB, but not in its absence. B. Cry2-mCherry-IBAR construct shows smaller decrease in cytosolic fluorescence intensity (relative to Cry2-mCherry) in the presence and absence of plasma membrane anchored CIB, indicative of significant pre-localization to plasma membrane. Blue arrows indicate beginning of blue light illumination (50 ms exposure of 480 nm light every 30 s); black arrows indicate end of blue light illumination. Graphs show changes in normalized cytosolic mCherry fluorescence.

Homo-oligomerization of the CRY-BAR constructs IBAR-Cry2-mCh and Cry2-mCh-IBAR results in membrane deformation and restriction of cellular dynamics (**Supporting Movies 1A and 1B**). These effects are reversible in the dark. To demonstrate this using Cry2-mCh-IBAR, we conducted a 10 min light activation sequence, followed by a 30 min observation period in the absence of blue light. Rapid restriction of membrane dynamics was observed during the 10 min blue light activation period, with full reversibility of membrane rounding achieved after 30 min in the absence of blue light (**Fig. 4; Supporting Movie 2**). We subsequently demonstrated that this effect can be selectively induced using localized illumination of the cell on a confocal microscope (**Fig. 5, Supporting Movie 3**). We also investigated whether the presence of a WH2 domain (present in IBAR-Cry2-mCh-WH2, but not in IBAR-Cry2-mCh and Cry2-mCh-IBAR) impacted the membrane remodeling capabilities of the optogenetic switch. Light activation of a N-terminal I-BAR domain without a C-terminal WH2 domain (IBAR-Cry2-mCh) promoted its accumulation in numerous filopodia-like protrusions that rapidly coalesced with continued light exposure (**Fig. 6A**). By contrast, light activation of a N-terminal I-BAR domain with a C-terminal WH2 domain, while not diminishing the overall light responsivity of the protein fusion (**Fig. 2A** and **2B**) inhibited pronounced filopodial accumulation (**Fig. 6B**). This result may illustrate competing functions of critical MTSS-1 protein domains, where the I-BAR domain is required for membrane binding, and the WH2 domain is required for recruitment of actin polymerizing components that promote protrusion formation and elongation (Saarikangas et al., 2011). Finally, a side-by-side comparison of light activation of IBAR-Cry2-mCh, IBAR-Cry2-mCh-WH2, and Cry2-mCh-IBAR demonstrated that more robust membrane remodeling activity is associated with light activation of Cry2-mCh-IBAR (**Supporting Movie 4**). This is likely due to enhanced PIP2 binding ability of the IBAR domain when in the C-terminal position of the optogenetic fusion protein.

**Figure 4.**
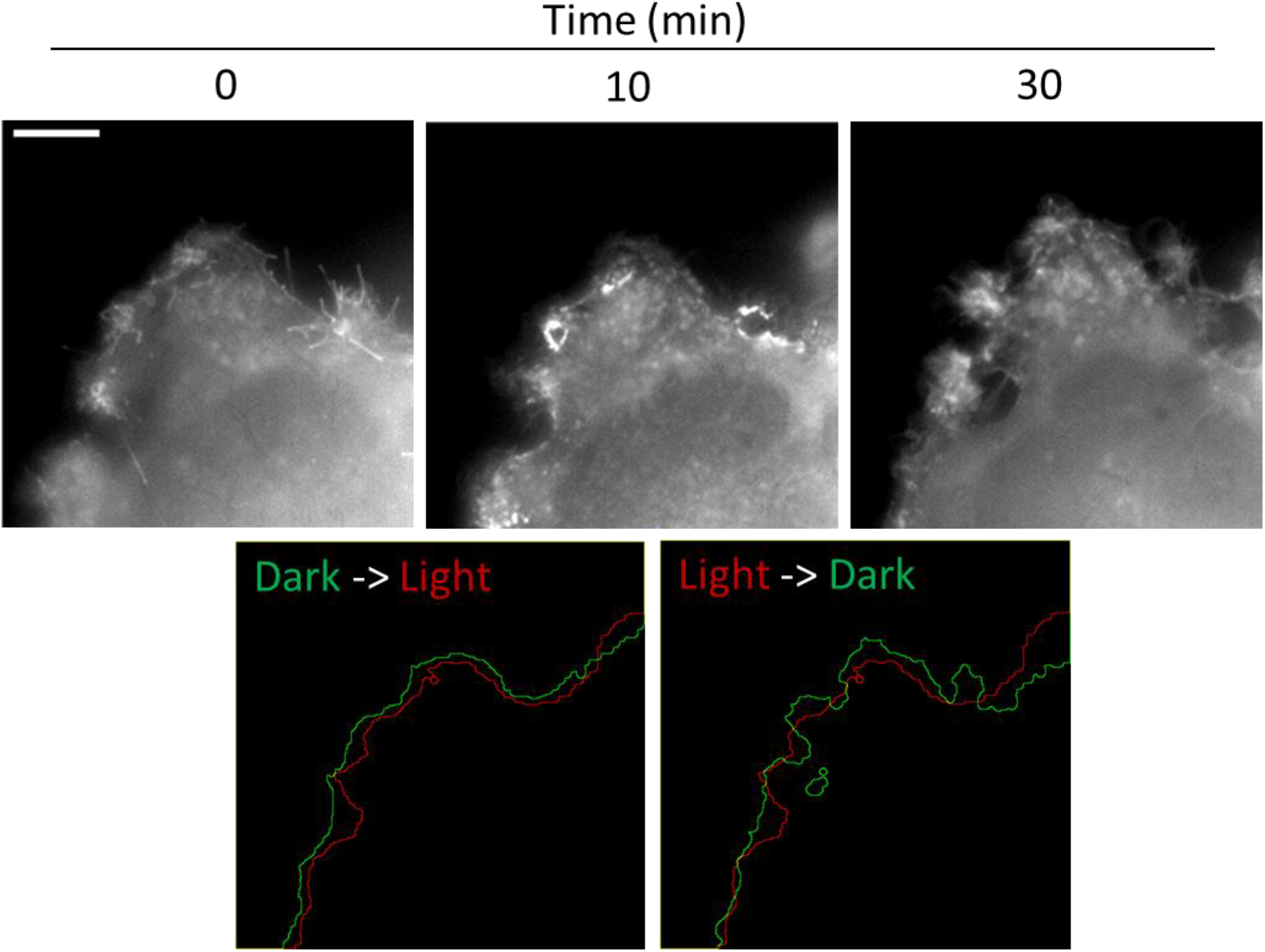
Reversible light-activated membrane remodeling with CRY-BAR. HEK293T cells transfected with Cry2-mCh-IBAR were subjected to blue light illumination for 10 minutes, followed by 30 minutes without blue light. Membrane retraction and reshaping is observed during blue light illumination, followed by recovery in the absence of blue light. Scale bar = 10 microns.

**Figure 5.**
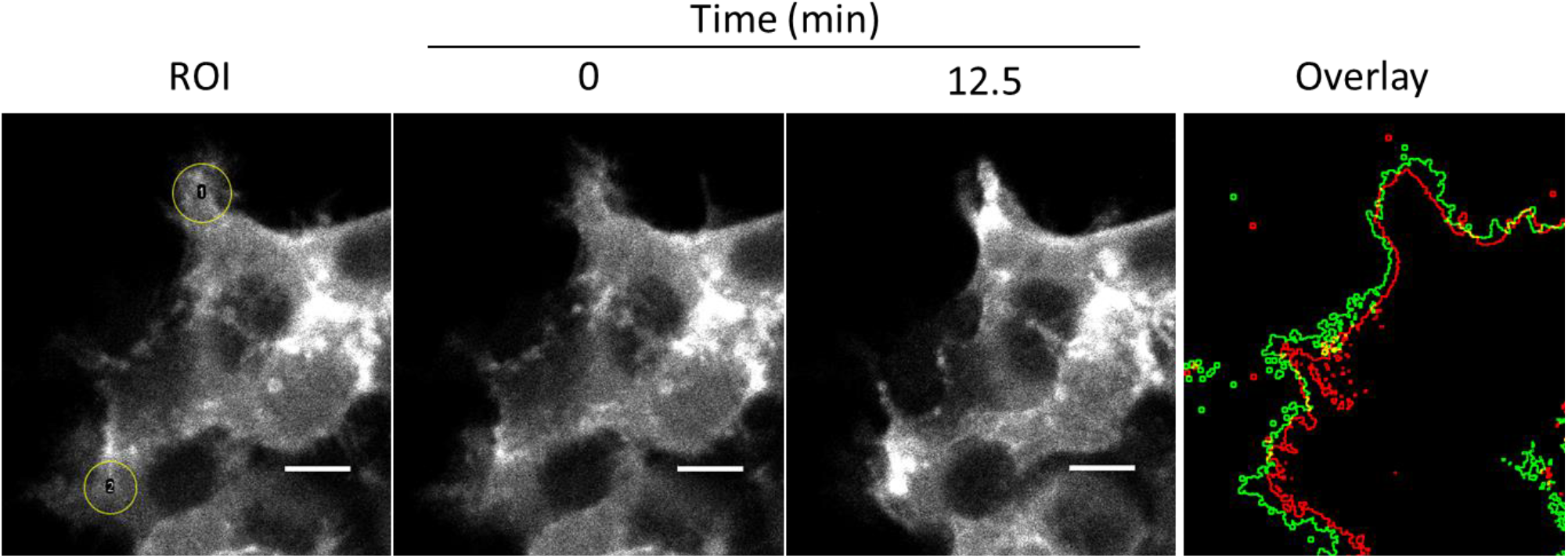
Localized light-activation of membrane remodeling with CRY-BAR. HEK293T cells transfected with Cry2P-mCh-IBAR were subjected to restricted blue light illumination (yellow circles) using a confocal microscope. Clustering of CRY-BAR is apparent in the areas irradiated with blue light. An overlay of the cell outline is shown before (green) and after (red) blue light illumination. Frames acquired every 30 s. Scale bar = 10 microns.

**Figure 6.**
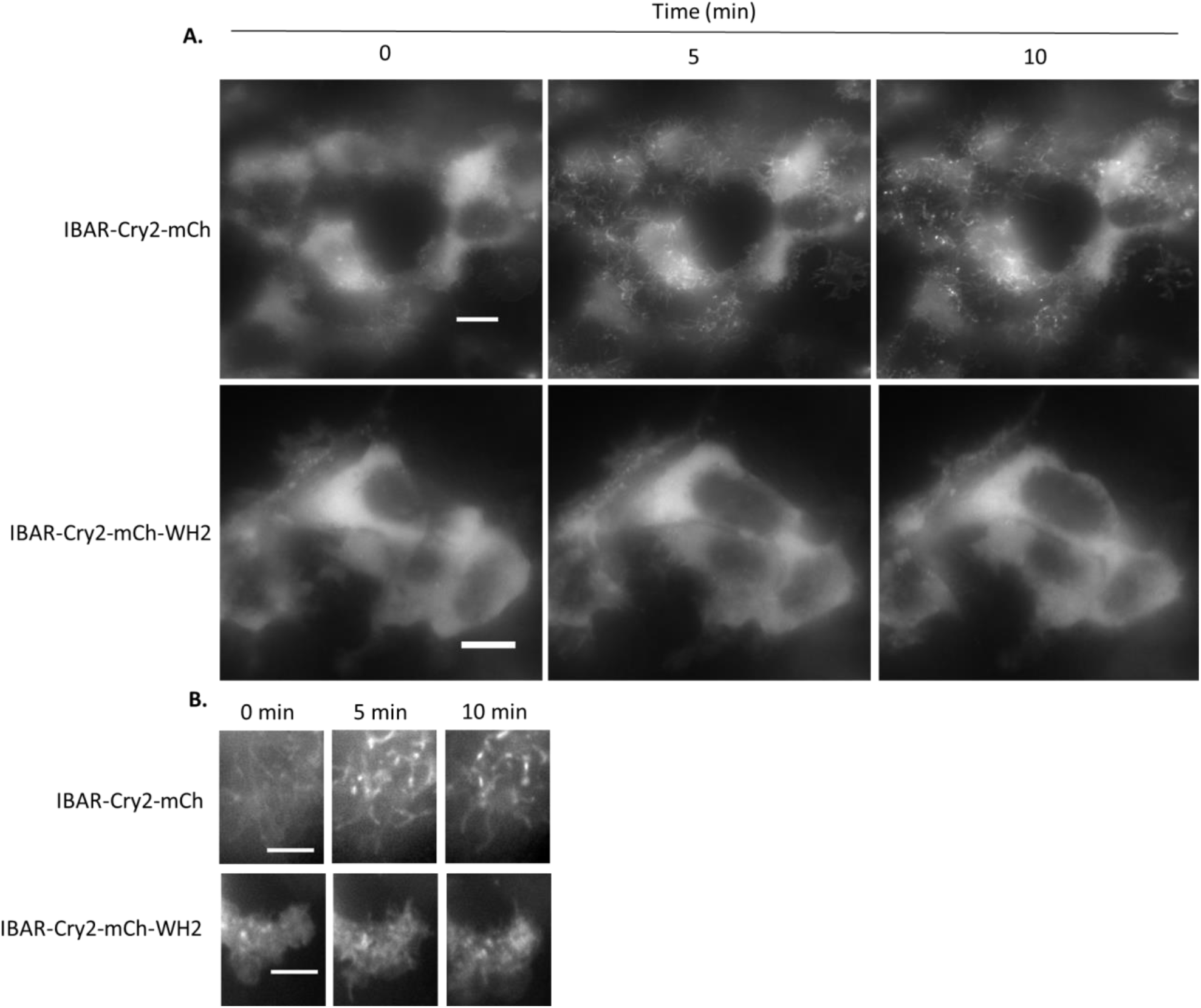
Light-activation of N-terminal IBAR-Cry2 fusions with and without a C-terminal WH2 domain. A. HEK293T cells transfected with IBAR-Cry2-mCh or IBAR-Cry2-mCh-WH2 were subjected to blue light illumination (50 ms pulse every 30 s) using a widefield microscope. IBAR-Cry2-mCh rapidly coalesces into dynamic filopodial structures. Scale bars = 10 microns. B. The presence of a WH2 domain inhibits accumulation into filopodial structures and subsequent coalescence. Scale bars = 5 microns.

We postulated that the dynamic membrane-remodeling activity observed with CRY-BARs might be due to its interaction with proteins that link the plasma membrane with the cytoskeleton. Ezrin is one such protein that has both lipid binding and cytoskeleton binding domains (Tsai et al., 2018). We hypothesized that clustering of the PIP2-bound CRY-BARs might impact dynamics by also clustering Ezrin, resulting in restriction of cytoskeletal-associated phenomenon such as membrane ruffling. To investigate this, we co-transfected Cry2-mCh-IBAR with an Ezrin-GFP fusion. In the presence of light, we observed co-localization of Ezrin-GFP with Cry2-mCh-IBAR clusters at the plasma membrane (**Fig. 7**). This effect was pronounced in the light, and absent in the dark, indicating that CRY-BAR activation actively restricts Ezrin dynamics. Cry-BAR activation accompanied by Ezrin sequestration also resulted in increased cell thickness (**Fig. 8**), with no such effect being observed in cells expressing the Cry2-mCh control.

**Figure 7.**
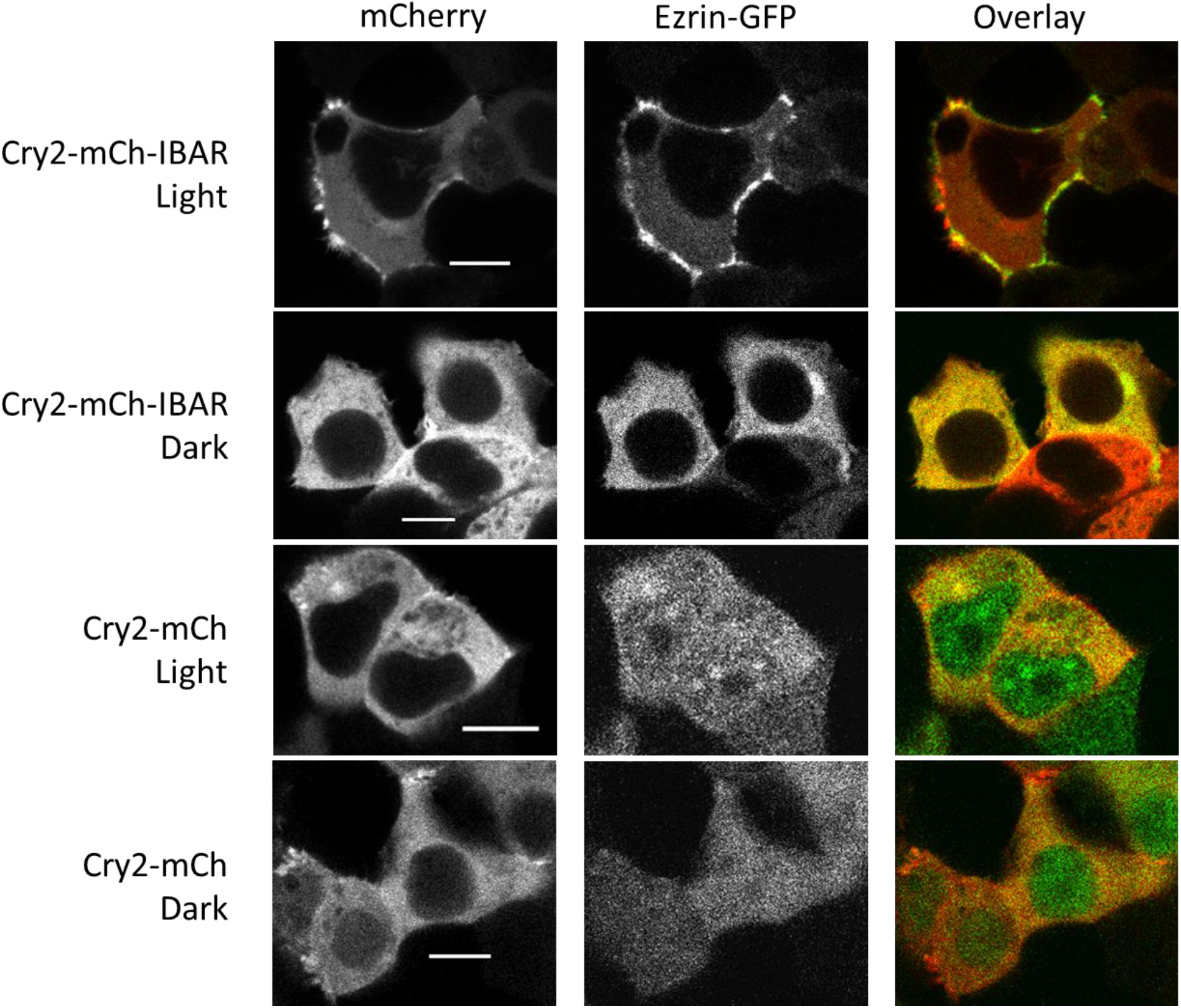
CRY-BAR activation linked to Ezrin. HEK293T cells co-transfected with Cry2-mCh-IBAR and Ezrin-GFP were subjected to blue light illumination or dark followed by fixation. Confocal microscopy reveals co-localization of CRY-BAR and Ezrin-GFP as a result of CRY-BAR activation. Scale bar = 10 microns.

**Figure 8.**
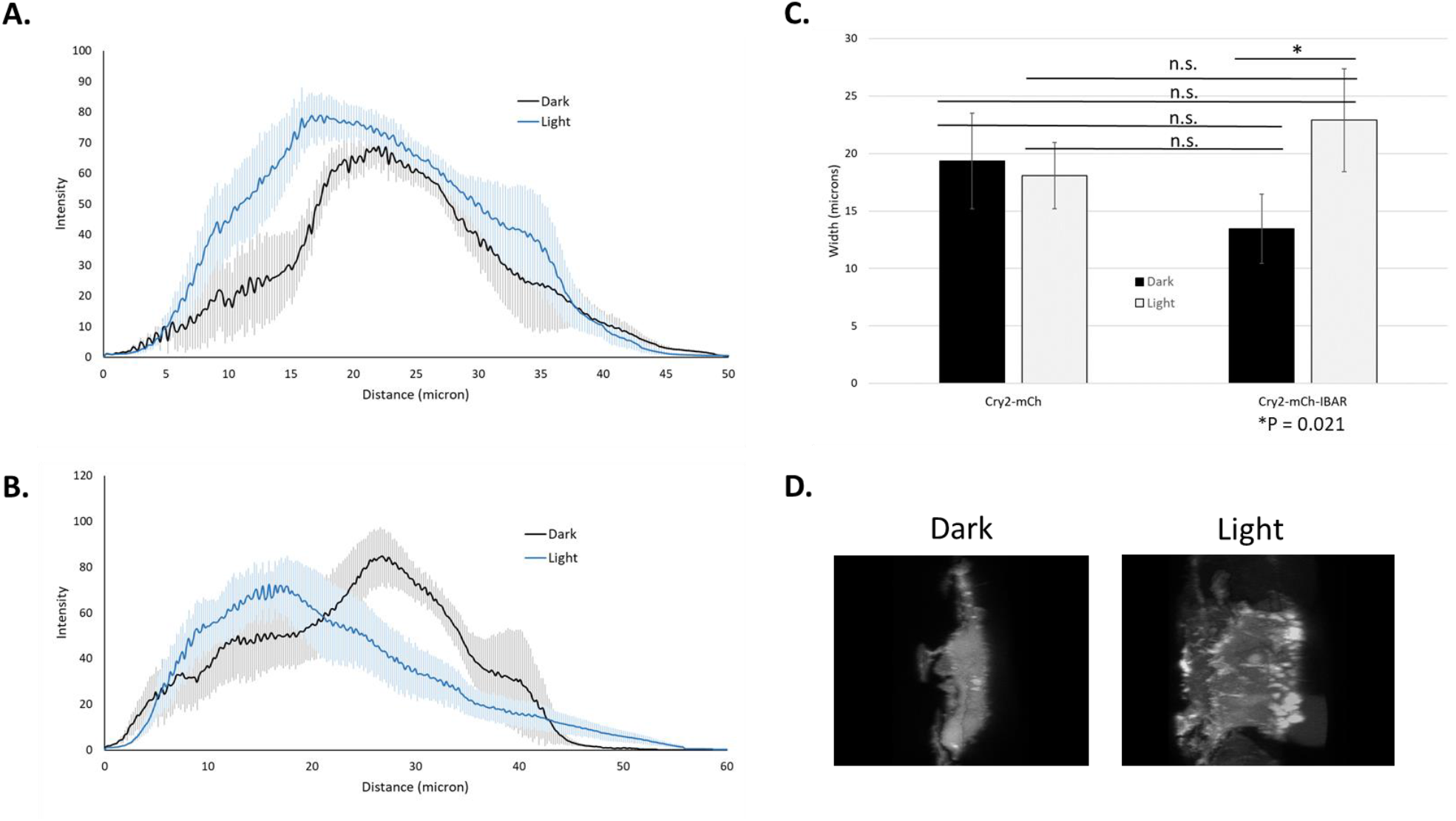
CRY-BAR activation and cell thickness. HEK293T cells co-transfected with Cry2-mCh-IBAR and Ezrin-GFP were subjected to blue light illumination or dark followed by fixation. Cellular thickness was analyzed for n=4 cells per experimental condition. A. Average cellular thickness of Cry2-mCherry-IBar transfected HEK293T cells before and after illumination. B. Average cellular thickness of Cry2-mCh transfected HEK293T cells before and after illumination. C. Analysis of average cellular thickness at half height (One way ANOVA: *p = 0.021; n.s.: p>0.05). D. Representative images of Z-projections of Ezrin-GFP distribution in Cry2-mCherry-Ibar transfected cells before and after illumination and fixation.

Having investigated the CRY-BAR response in immortalized cell culture, we next evaluated the capability of the sensor to report membrane expansion dynamics in neurons. For this, dissociated postnatal cortical neuron cultures prepared from newborn mice were transfected with Cry2-mCh-IBAR, or co-transfected with CIB-GFP-CAAX, according to protocol we developed for expressing optogenetic sensors in primary cultures (Bunner et al., 2021). Light activation of Cry2-mCh-IBAR, but not Cry2-mCh, resulted in elongation of neuronal processes that might indicate early stage spinogenesis (**Fig. 9, Supporting Movies 5 and 6**). There was no apparent benefit to activation of Cry2-mCh-IBAR in the presence of CIB-GFP-CAAX, providing additional evidence that activation of Cry2-mCh-IBAR alone is sufficient for effecting membrane dynamics (**Fig. 9C**). Accordingly, whole cell illumination of neurons transfected with only Cry2-mCh-IBAR results in abundant membranous cluster formation throughout neuronal processes (**Supporting Figure 2**). Finally, using localized illumination conditions, we demonstrated that Cry2-mCh-IBAR can be selectively activated in neuronal processes (**Fig. 10, Supporting Movie 7**). Taken together, this work provides evidence that the Cry2-mCh-IBAR molecular optogenetic tool is suitable for the targeted manipulation of cytoskeletal structures and dynamics at the plasma membrane.

**Figure 9.**
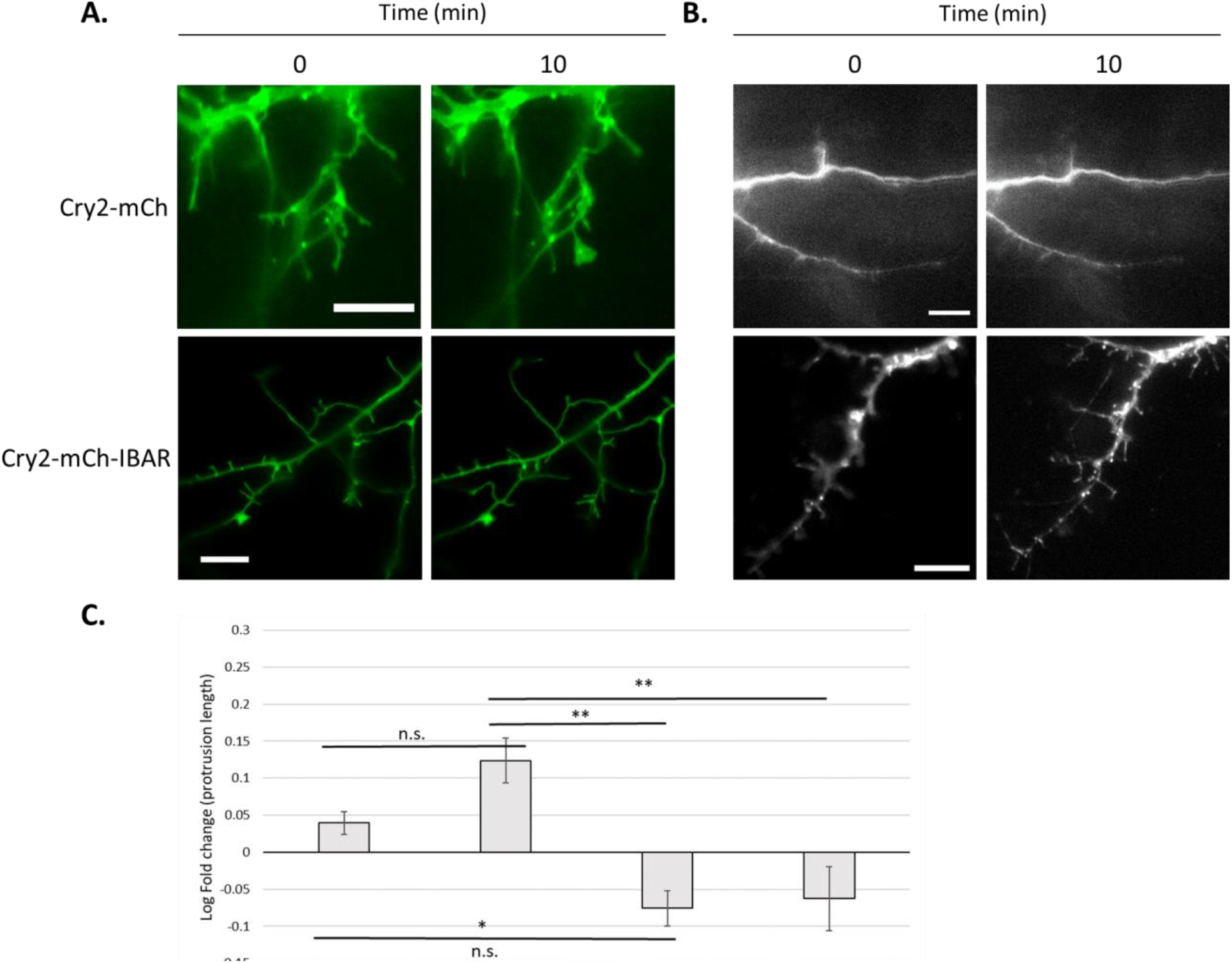
CRY-BAR plasma membrane recruitment in neurons. Pre- and 10 min-post 480 nm light illumination of processes in hippocampal neurons: A. co-transfected with Cry2-mCherry-IBAR or Cry2-mCh and CIB-GFP-CAAX (GFP channel shown) and B. transfected with Cry2-mCherry-IBAR or Cry2-mCh. C. Log fold-increase in protrusion length in protrusions of Cry2-mCh-IBAR and Cry2-mCh expressing neurons with or without CIB-GFP-CAAX after 10 minutes of 480 nm light illumination (n = 10 neurons per condition; One-way ANOVA: **p<0.001; *p=0.04; n.s.: p>0.05).

**Figure 10.**
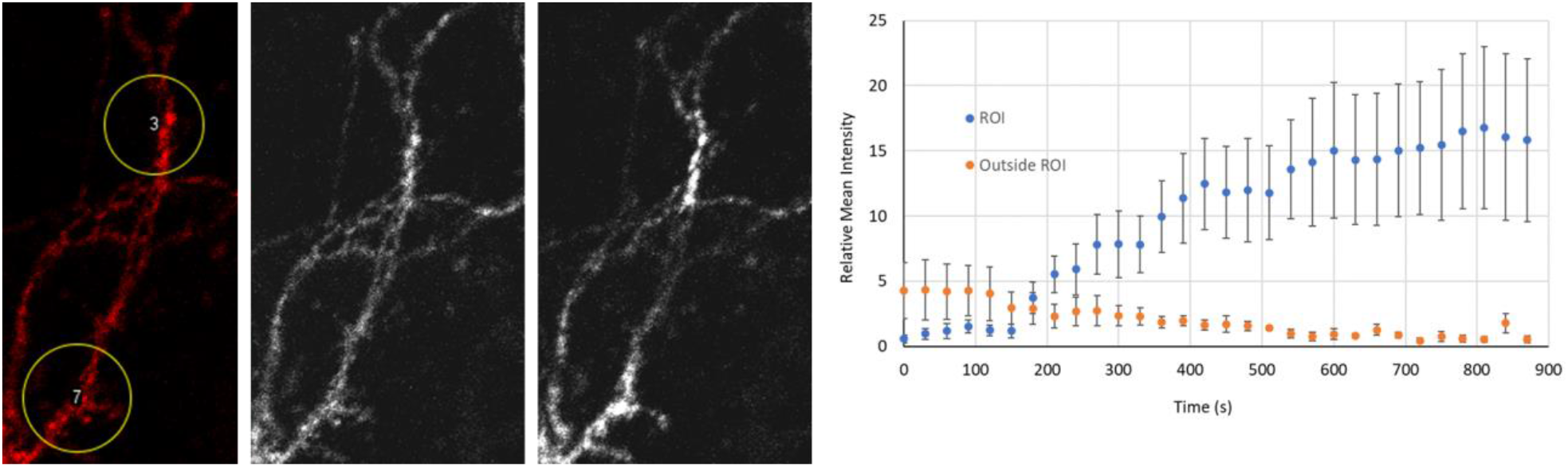
Localized light-activation of CRY-BAR in primary neurons. Neurons transfected with Cry2-mCh-IBAR were subjected to restricted blue light illumination (yellow circles) using a confocal microscope. Clustering of CRY-BAR is apparent in the areas illuminated 15 min post-blue light stimulation. A plot of fluorescence intensity from four illuminated ROIs vs. four non-illuminated regions is shown at right (error bars = SEM). Scale bar = 10 microns.

## Conclusion

In this report, we describe the development of a family of optogenetic switches, collectively named Cry-BAR, that comprise a versatile platform for controlling membrane dynamics in live cells with high spatial and temporal resolution. These switches combine homo- and hetero-oligomerization of the Cry2/CIB photoreceptor system with the innate PIP2 binding affinity of the I-BAR domain. As their function varies depending on the presence of various functional elements, such as WH2 binding domains, we anticipate that a modular approach can be undertaken to adapt these optogenetic switches for other applications, including the recruitment of enzyme and receptor activating and inhibitory domains. Finally, Cry-BARs are suitable tools to study membrane dynamics not only in immortalized cells, but also in sensitive and difficult to transfect primary cultures, such as neurons. Therefore, the Cry-BAR optogenetic switches we report here are expected to have wide applicability for investigating cellular processes associated with membrane dynamics in a variety of experimental paradigms.

## Authors’ contributions

AIW, WPB, ESM, and RMH designed experiments. AIW and RMH performed cloning and cell culture experiments. WBP and ESM prepared neuron cultures and transfected neurons. AIW, WPB, ESM, and RMH analyzed data and prepared figures. All authors contributed to writing the manuscript and agreed to the content of this paper.

## Acknowledgements

Authors wish to thank Mr. Collin T. O’Bryant for assistance with confocal microscopy. This work was supported by a seed grant to RMH and ESM from the East Carolina University Biomaterials Research Cluster.

## Availability of Materials and Data

Experimental data are available upon request. All Cry-BAR plasmids will be available through Addgene (Watertown, MA), a nonprofit DNA repository.

---

**Supporting Figure1.**
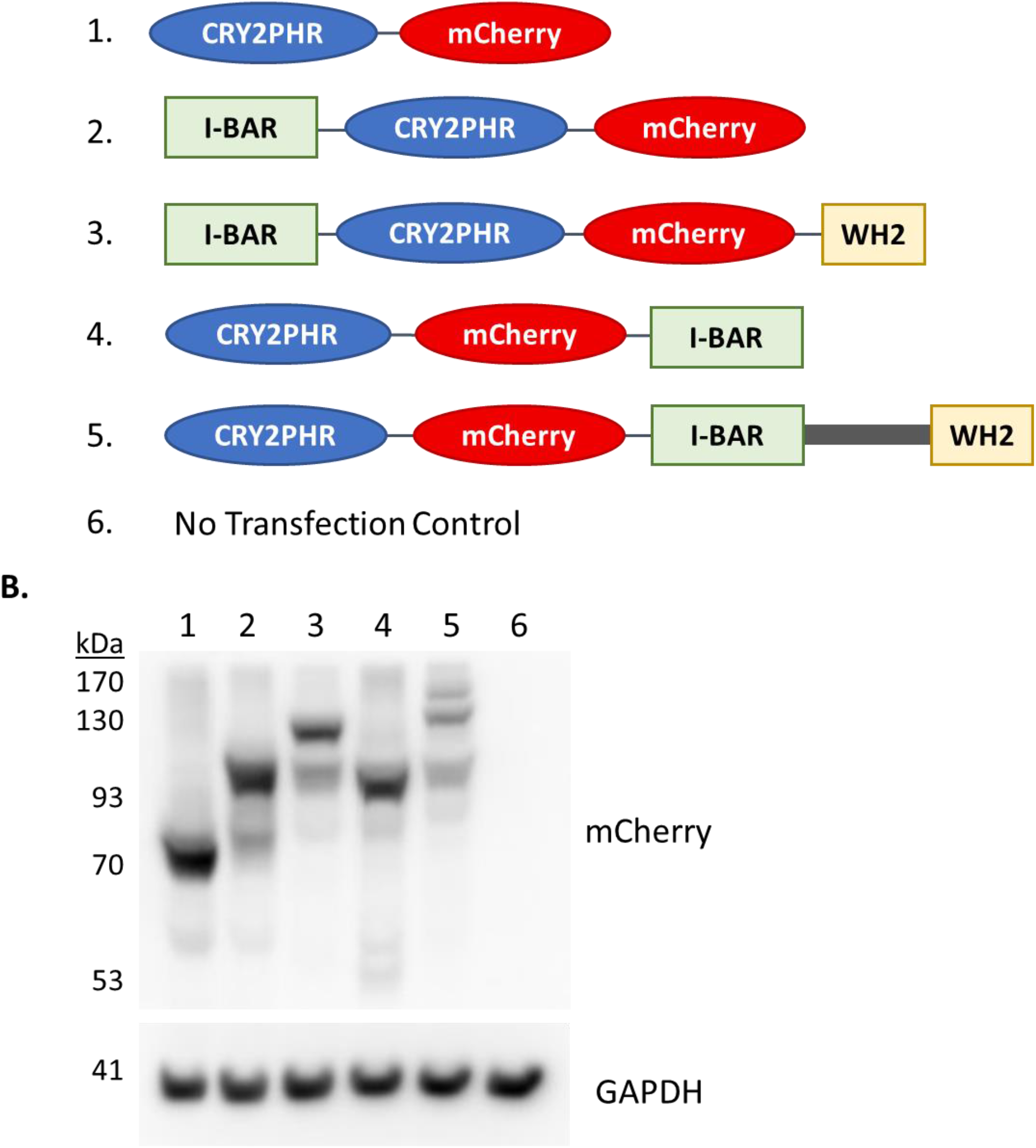
CRY-BAR construct design and expression. A. Diagrams of the CRY-BAR protein fusions; Constructs 2, 3, and 4 contain portions of the MTSS-1 protein (I-BAR, WH2, or both); Construct 5 contains the full length MTSS-1 protein. B. Lysates of HEK293T cells transfected with CRY-BAR constructs and controls (Lanes 1 – 6, numbers correspond to panel A) were western blotted with anti-mCherry antibody. Expected Molecular Weights: 1. 85 kD; 2. 111 kD; 3. 130 kD; 4. 111 kD; 5. 168 kD; 6. No transfect.

**Supporting Figure2.**
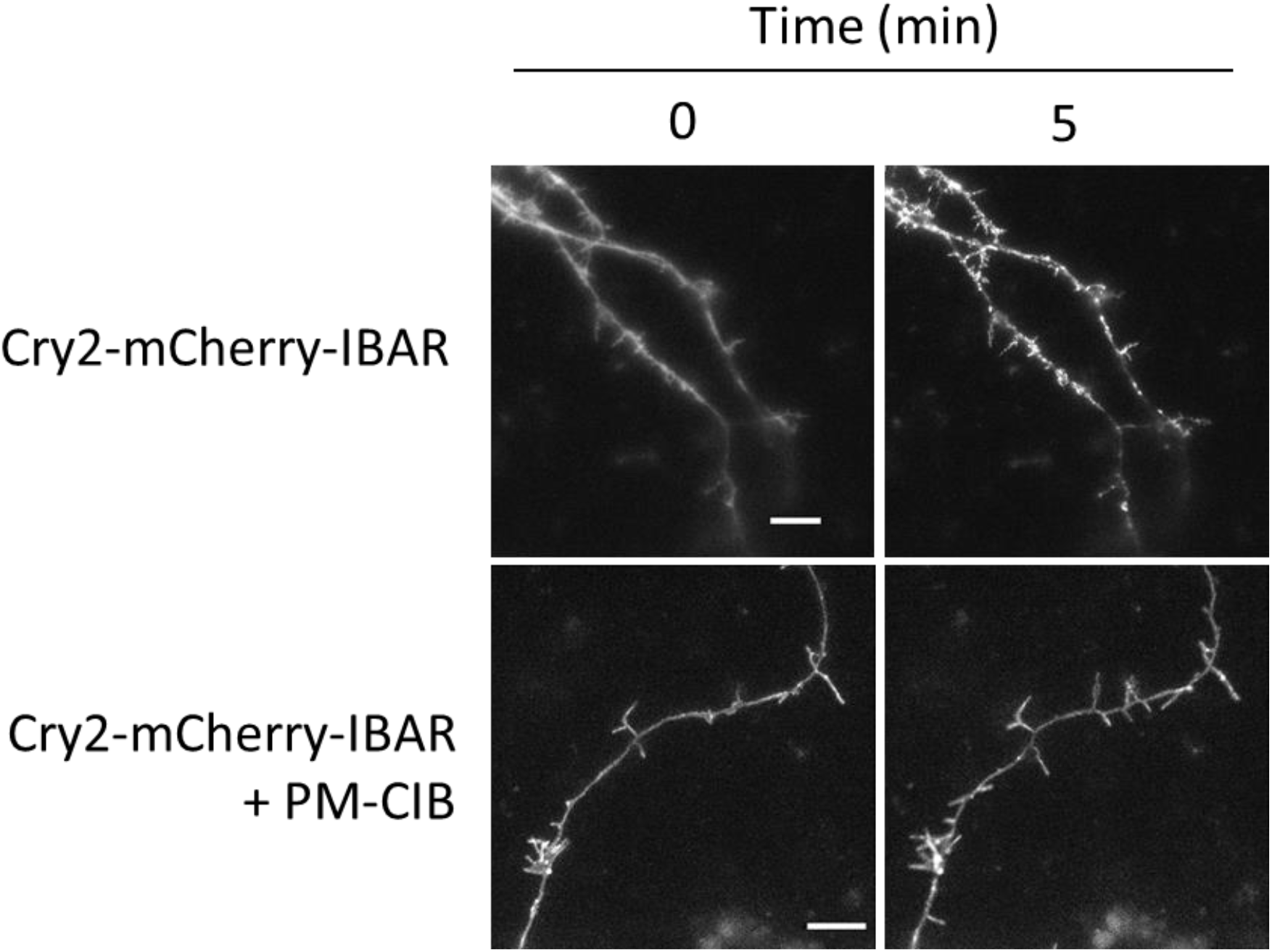
Global activation of CRY-BAR in hippocampal neurons. Neurons transfected with Cry2-mCh-IBAR; and co-transfected with Cry2-mCh-IBAR + PM-CIB were illuminated every thirty seconds with 480 nm light on a widefield microscope. After 5 minutes, Cry2-mCh-IBAR forms numerous clusters throughout the neuronal processes, whereas in the presence of PM-CIB no such clusters are formed. Scale bar = 10 microns.

## Supporting Movie Legends

**Supporting Movie 1A** Cry2-mCh-IBAR light activation in HEK293T cells

**Supporting Movie 1B** Cry2-mCh-IBAR light activation in HEK293T cells

**Supporting Movie 2** Cry2-mCh-IBAR light activation in HEK293T cells – zoomed in region

**Supporting Movie 3** Localized Cry2-mCh-IBAR light activation in HEK293T cells

**Supporting Movie 4** Side-by-side light activation of CRY-BARs in HEK293T cells

**Supporting Movie 5** Light activation of Cry2-mCh-IBAR/CIB-CAAX in neuronal process

**Supporting Movie 6** Light activation of Cry2-mCh-IBAR in neuronal processes

**Supporting Movie 7** Localized Cry2-mCh-IBAR light activation in neuronal process

