## Supplementary figures and images for "CRY-BARs: Versatile light-gated molecular tools for the remodeling of membrane architectures"

### Supporting Movie 1A

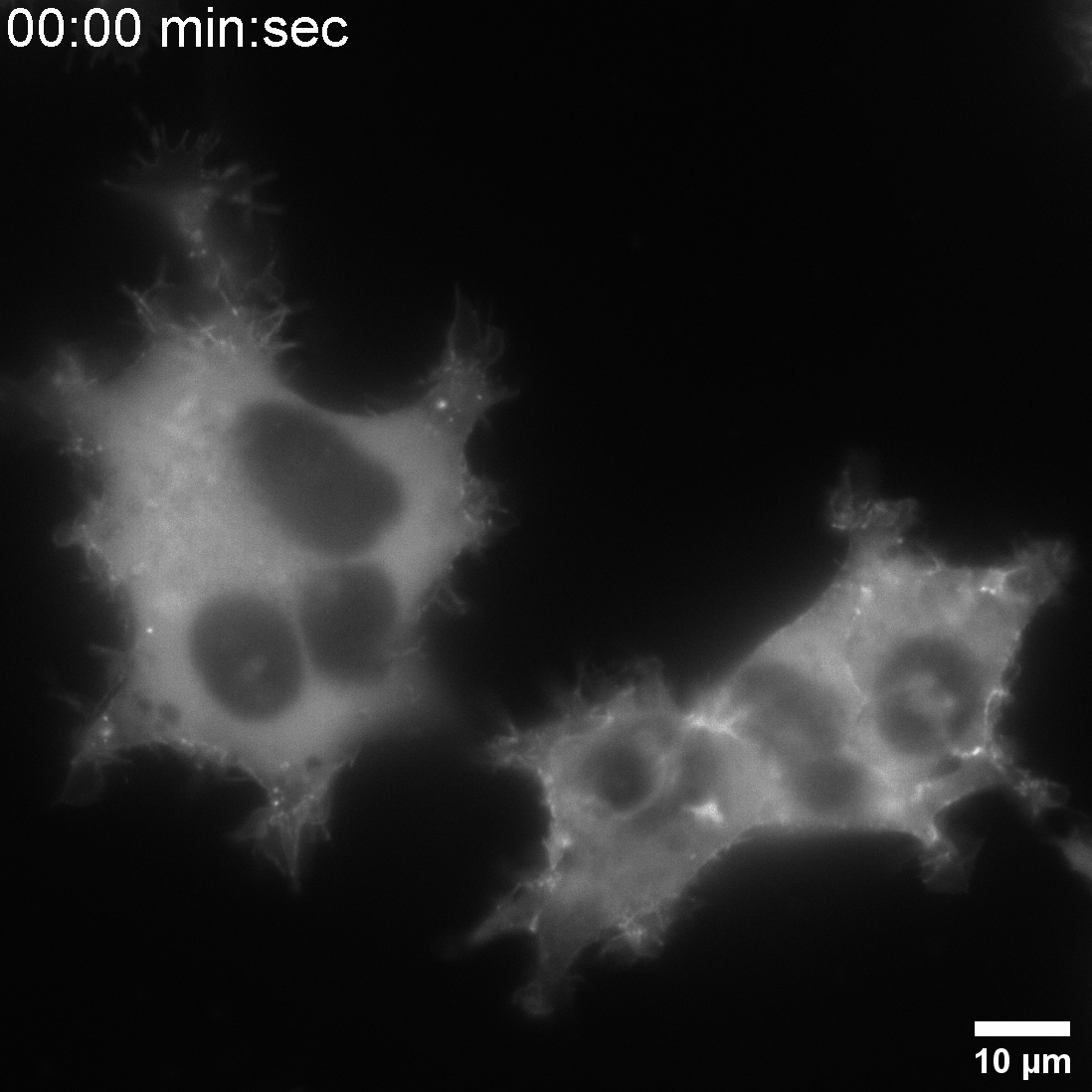

### Supporting Movie 1B

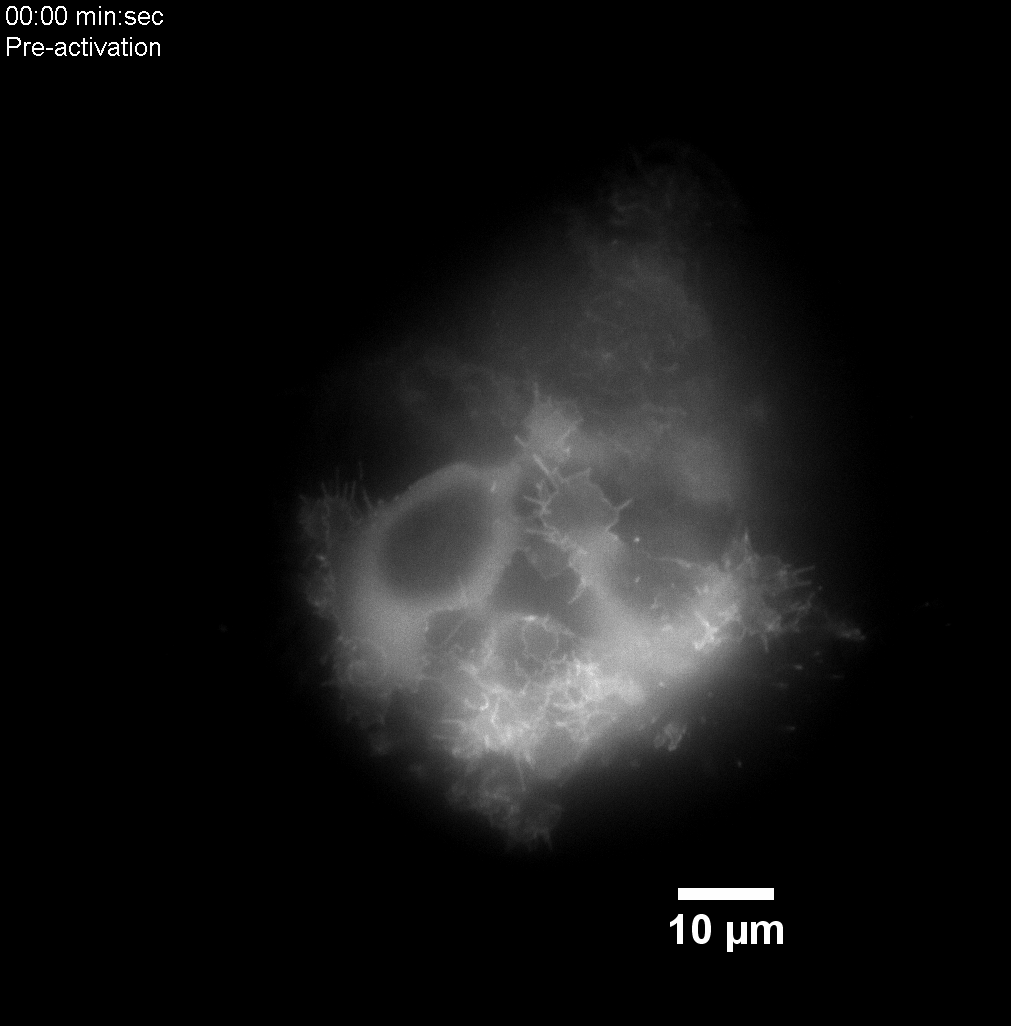

### Supporting Movie 2

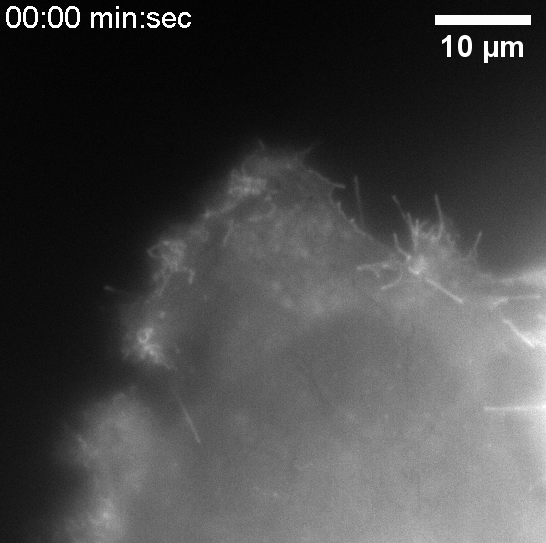

### Supporting Movie 3

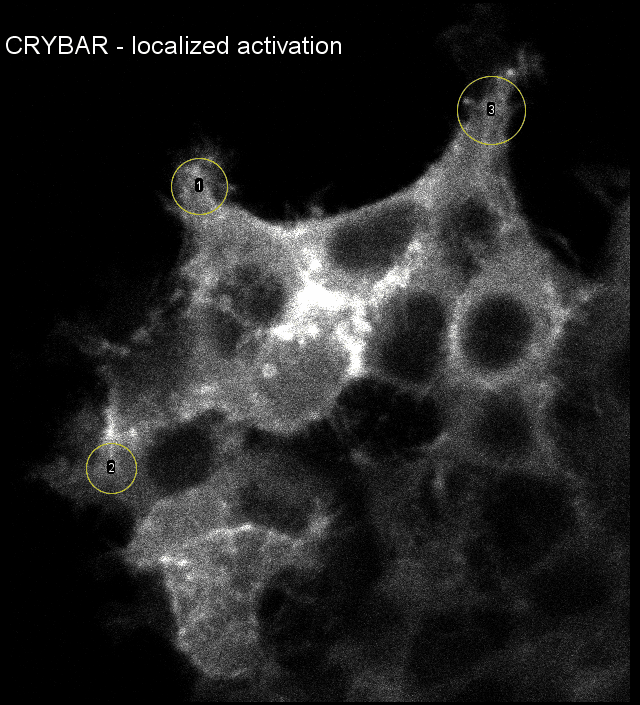

### Supporting Movie 5

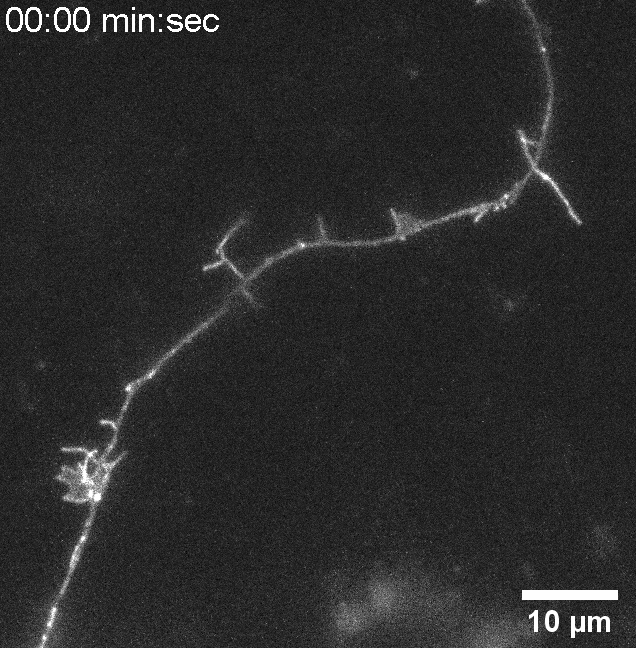

### Supporting Movie 6

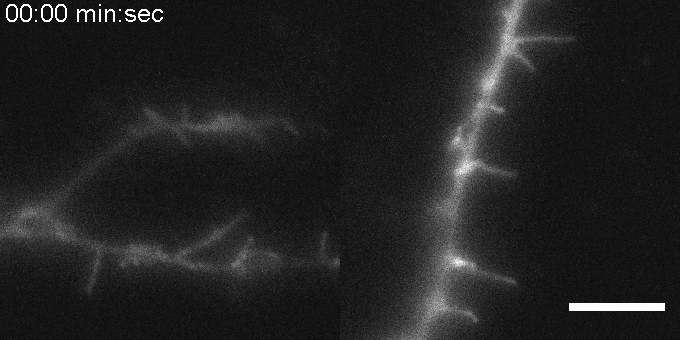

### Supporting Movie 7

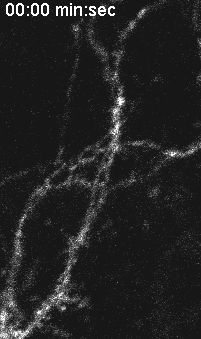
